# predicTox: An integrated database of clinical risk frequencies and human gene expression signatures for cardiotoxic drugs

**DOI:** 10.1101/2025.05.23.655401

**Authors:** Jens Hansen, Pedro Martinez, Arjun S. Yadaw, Yuguang Xiong, Rebecca Racz, Michael R. Pacanowski, Laura L. Hopkins, Nicholas M. P. King, Darrell Abernethy, Eric Sobie, Ravi Iyengar

## Abstract

We have used drug-induced transcriptomic responses and whole-genome sequences in healthy human induced pluripotent stem cells (iPSC)-derived cardiomyocyte lines to identify cellular functional pathways and genomic variants potentially associated with the cardiotoxic effects of tyrosine kinase inhibitors (TKIs) and other cancer drugs. Here, we describe predicTox.org, an interactive website that organizes our data and its integration with knowledge from cell pathways and genomic databases. DrugTox summary cards give results of these analyses and metadata for each drug. Fields include FDA Adverse Event Reporting System (FAERS) cardiotoxicity risk scores, cell pathways, and genomic variants potentially associated with drug-induced cardiotoxicity. At a detailed level, predicTox provides ranked lists of pathway signatures that are common for cardiotoxic TKIs, up- and downregulated pathways associated with cardiotoxicity induced by specific TKIs and TKI-regulated genes mapping to those pathways. predicTox provides downloadable lists of drug-induced differentially expressed genes (DEGs) and pathways, drug-related genomic variants associated with cardiotoxicity, all with their statistical metrics, and mathematical models to simulate drug effects on heart physiology. Building on the results of our algorithm for independent reidentification of the well-known *rs2229774* variant for anthracycline-induced cardiotoxicity, we describe how our data can be queried to identify potential variants associated with drug-induced cardiotoxicity by affecting a drug’s pharmacodynamics and pharmacokinetics.

## Introduction

Adverse drug reactions (ADR) continue to be a major driver of failure in drug development 1 and adverse drug reactions may go undetected until a drug has been in clinical use for several years. Methods to characterize the mechanisms of ADRs and predict their occurrence are critical to improving patient safety. To close this gap, several integrative approaches have been proposed over the past decades. These include the FDA’s critical path initiative ^2^ and efforts at developing predictive safety testing consortia. ^3^ Abernethy and his colleagues at the FDA proposed the use of systems biology methods to develop predictive models for drug safety.^4^ The predicTox Knowledge Environment (predicTox - KE), a concept first articulated by Abernethy, was an outgrowth of these efforts.

Cardiotoxicity whether affecting cardiac structure or function, is recognized as a major complication of cancer therapies, including tyrosine kinase inhibitors.^5-7^ Human iPSC derived-cardiomyocytes have emerged as a useful model system to study cardiac diseases ^8^ and drug action including adverse events.^9^ These human in-vitro assay systems can also be used to identify the potential for adverse events associated with therapeutic drugs. This information, in the form of signatures that are ranked lists of genes or pathways altered upon drug treatment, can be used to predict the potential toxicity of new drug candidates from cell-based assays. The predictions can be useful in early cell-based safety studies during drug development to minimize the use of animal models for drug safety testing and ascertain relevant information from humans early in development. Since iPSCs from individual subjects are used and when genomic data are available, these studies can also be used to predict if a human subject may be susceptible to drug toxicity based on genomic variants affecting the pharmacokinetics (PK) and pharmacodynamics (PD) or affecting the pathway genes perturbed by drug treatment.

We have previously used drug-induced gene expression studies in human heart cells ^10^ and iPSC-derived cardiomyocytes ^11^ to obtain transcriptomic signatures associated with FDA-approved drugs with evidence of cardiotoxicity. To create a resource for researchers investigating new compounds or factors that may contribute to cardiotoxicity, we created a knowledge environment. The predicTox-KE is a database that has organized the drug-induced gene expression data in cardiomyocytes derived from iPSCs of healthy human subjects and integrated the transcriptomic data with clinical adverse event data, and human genomic variant data in a manner that can be widely applied. The predicTox-KE provides gene- and cell-level regulatory pathway information from human iPSC-derived cardiomyocytes.

## Materials and Methods

We provide a summary of our published algorithms that we used to generate the data presented on the predicTox website. For details and flow charts please see our research publication.^11^ Code for reproduction can be downloaded from GitHub.^12^

### Adverse Event Estimation

Using data from FAERS,^13^ we calculated the reporting Odds Ratio (ROR) and 95% confidence interval for each drug to be associated with the cardiac disorder AE (OAE 0000084) as defined by the Ontology of Adverse Events (OAE).^14^ Drugs were ranked by ROR. A downloadable PowerPoint presentation that can be accessed from the “Drugs” page provides details on used algorithms.

### Signatures of ranked differentially expressed genes

Differentially expressed genes (DEGs) induced by each drug in each cell line were identified using edgeR.^15^ Pairwise correlation analysis followed by hierarchical clustering grouped DEGs mostly by treated cell line and not by administered drug. This indicates that the transcriptional responses are dominated by cell-line-selective effects that hide drug-selective responses. To reveal the drug-selective response vectors, we applied singular value decomposition (SVD) to identify drug-selective subspaces that increase the similarity of the projected transcriptional response vectors for the drug of interest. Among potential drug-selective subspaces, we prioritized subspaces that detect a transcriptomic outlier response in one cell line and similar responses in the other cell lines. If this was not the case, we emphasized similarity over information loss relative to the complete responses. Gene expression vectors projected onto each selected drug-selective subspace were defined as the corresponding transcriptional responses. Drug-selective DEGs were averaged across all cell lines.

### Pathways predicted to be associated with cardiotoxicity

Up- and downregulated genes among the top 600 drug-selective DEGs within each cell line were subjected to pathway enrichment analysis using the Molecular Biology of the Cell Ontology (MBCO).^16,17^ Predicted (MBCO level-3) pathways were ranked by significance. For each rank from 1 to 30, we counted how many cardiotoxic and non-cardiotoxic TKIs up- or downregulated a pathway of interest with the same or higher rank (lower rank number). This allowed calculation of precision, recall and F1 score at each rank position. For the F1 score, we emphasized the precision over recall (beta=0.25) to focus on pathways that are sensitive for cardiotoxicity. To prevent focus on an arbitrarily selected significance rank, we calculated the F1 score area under the curve (AUC) from ranks 1 to 30. Additionally, we subtracted 50% of the AUC for a downregulated pathway from the AUC of an upregulated pathway and vice versa to prevent conflicting results that would be hard to interpret. Final AUCs for up- and downregulated pathways were combined and ranked (AUC rank). The top 25 pathways are presented.

### Genomic variants potentially associated with cardiotoxicity

To influence a drug’s cardiotoxicity in a monogenic way a variant could map to a gene that participates in a pathway whose altered expression is associated with cardiotoxicity or map to a gene whose gene product participates in a drug’s pharmacokinetics (PK) or pharmacodynamics (PD). To meet common frequencies of drug-induced cardiotoxicity ^7^ in both cases, we considered only variants that contain alleles with population-wide frequencies of maximal 10%. Relevance for heart tissue was assumed for variants that are within exon-coding regions or are part of cis-expression-(cis-e-) or -splicing-quantitative trait loci (s-QTLs) in the heart ^18^. Pathway associated variants were identified by searching for variants that map to genes annotated to the pathways that we predicted to be associated with TKI-induced cardiotoxicity. Since the effect of a genomic variant on a drug’s PK/PD response occurs before the drug-induced transcriptional response, it should generate an outlier response. Based on statistical likelihood, either one or none of our cell lines should contain alleles within a variant that are associated with cardiotoxicity (or alternatively, cardioprotection). Consequently, we searched for single cell lines that show a transcriptional outlier response to a drug of interest. Within identified cell lines we searched for variant alleles that fulfill our population-wide and relevance requirements, have higher counts in the outlier cell line than in all other five cell lines and map to genes involved in that drug’s PK/PD mechanisms as curated from DrugBank.^19^

## Results and Discussion

predicTox-KE currently has signatures for a set of 54 drugs, including 23 small molecule tyrosine kinase inhibitors and four monoclonal antibodies against tyrosine kinases. The drugs were tested on human iPSC-derived cardiomyocytes from six healthy subjects of both sexes.^20^ Signatures were predicted from bulk transcriptomic studies. For users who want to identify cardiotoxicity potential for new drug candidates we provide summary signatures that are specifically associated with cardiotoxic TKIs. Cardiotoxicity as clinically assessed was obtained from the literature ^7^ and quantified by analyses of FDA Adverse Event Reporting System (FAERS) data.^13^ We highlight a few features that PredicTox-KE provides:

### Summary of Drug Action

For pharmacologists focused on drug action, we provide DrugTox Summary Cards that for each drug tested list cardiotoxic status and rank, top ranked genes and pathways induced by the drug as well as those induced pathways and genomic variants that are predicted to be associated with TKI cardiotoxicity. We provide a search function to identify all drugs that regulate a human gene of interest. In addition to the processed data, we provide all data sets for downloading and further analyses by the user.

### Genomic variants

For researchers interested in exploring the genomic underpinning of drug safety issues, predicTox-KE provide lists of genomic variants that map to PK/PD mechanisms, as well as variants associated with genes in functional pathways within cardiomyocytes predicted to be associated with TKI cardiotoxicity. These genomic variants could be used to identify human subjects who could potentially have cardiotoxic responses to drug therapy.

### DYNAMICAL models

The effects of drug action on cellular pathways are best understood as changes in time-dependent physiological responses. These are best captured by numerical simulations using differential equation-based models. predicTox provides downloadable MATLAB code for an arrhythmia model based on publication by Shim et al ^21^ and for genes involved in cardiomyocyte hypertrophy,^22^ a pathological response to cardiotoxic drugs. Both models for numerical simulations have Readme files and associated glossaries.

## Layout of the predicTox-KE

### Landing-Page

The landing page of the predicTox-KE provides an easy way to use the provided resources. In the header of the homepage we provide links to About, How-to-Use, and Glossary pages that help the user navigate the site. A search function allows the user to enter queries to search the list of 54 drugs for which transcriptomic signatures are available. Selecting one of the popup drug names opens the DrugTox summary card that provides an integrated view of all the cardiotoxicity data for the drug of interest. The card is described in detail below. The user can also search for gene names (i.e., human NCBI gene symbols) to get a summary of the drugs that up or down regulate the gene of interest in the cardiomyocyte cell lines.

The window “Summary of Signatures for Cardiotoxicity” below the search function directs the user to a page showing all predicted cardiotoxicity-associated pathways and the expressed pathway genes that we discuss below. The box “Downloadable Datasets” gives the user access to a range of unprocessed and processed data that we used for our analyses. These are organized in four sections: “Drugs”, “Adverse Events - Cardiotoxicity”, “Bulk Transcriptomic Datasets - Metadata” and “Models”. The layout of the home page is shown in Figure 1.

**Fig. 1.**
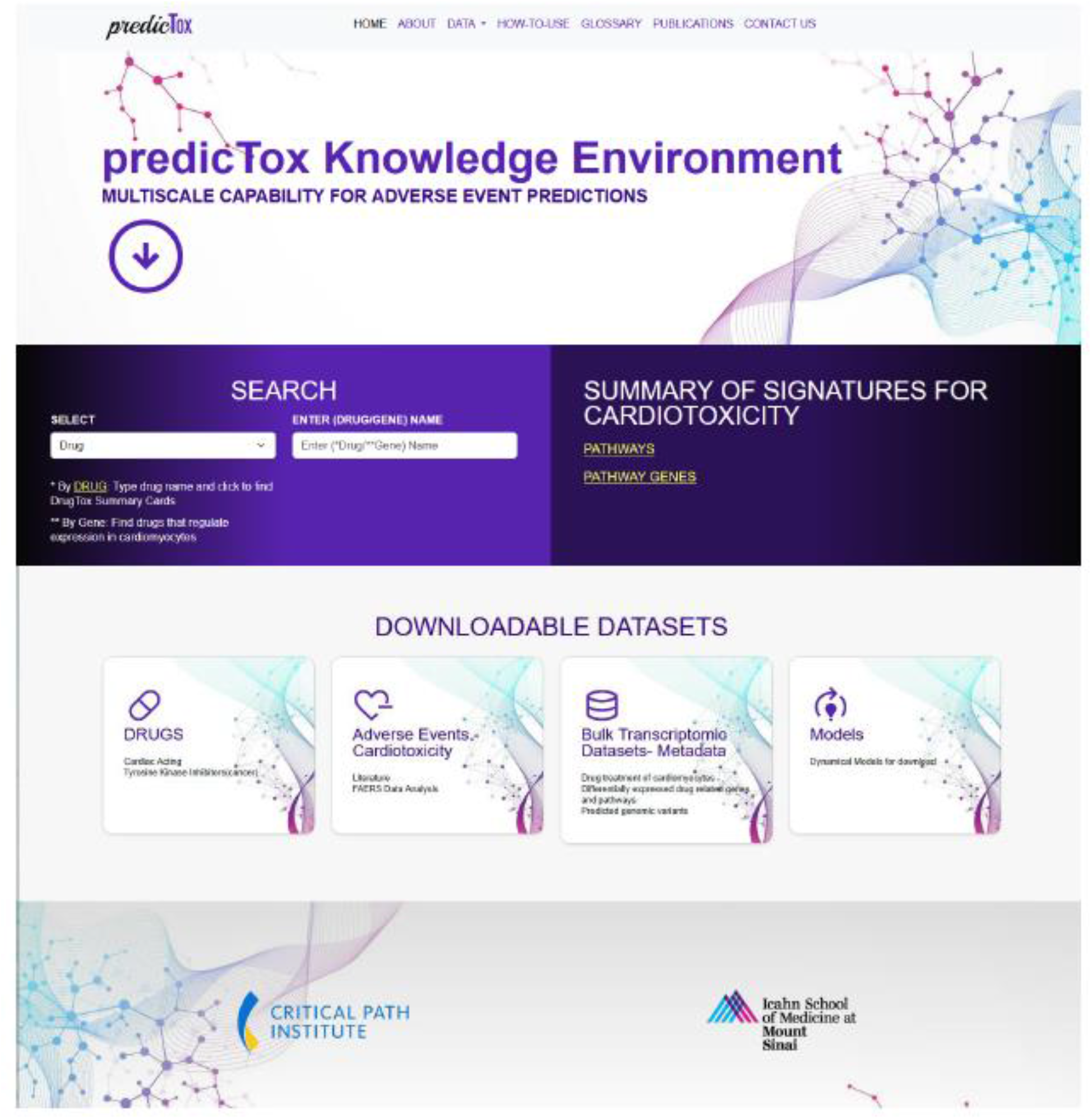
predicTox homepage. A screen shot of the home page which provides searchable DrugTox Summary cards and gene Summary Cards. The home page also provides clickable links to various pages that lists data sets and other downloadable materials. ND: not determined.

### DrugTox Summary Card

Selecting “Drug” in the “Select” list-box and entering a drug name into the “Enter (Drug/Gene) Name” textbox within the “Search” window, followed by clicking on the full name that pops up below opens a summary card that contains information in seven different domains (Fig. 2). The information provided on the card includes the top ranked genes and pathways that are affected by drug treatment as well as the number of drug-regulated pathways that are predicted to be associated with cardiotoxicity if the selected drug is cardiotoxic. The top three of the latter up- and downregulated pathways are given below. Additionally, for cardiotoxic drugs we provide the number of genomic variants that could harbor alleles associated with an increased (or decreased) frequency of cardiotoxicity. These constitute variants mapping to genes involved in pharmacodynamics (PD) or pharmacokinetics (PK) of the selected drug and to pathways predicted to be associated with cardiotoxicity and induced by the drug. The drug summary card can serve as a handy reference for users who may want to compare their experimental results with a drug candidate with the more summarized results presented here.

**Fig. 2.**
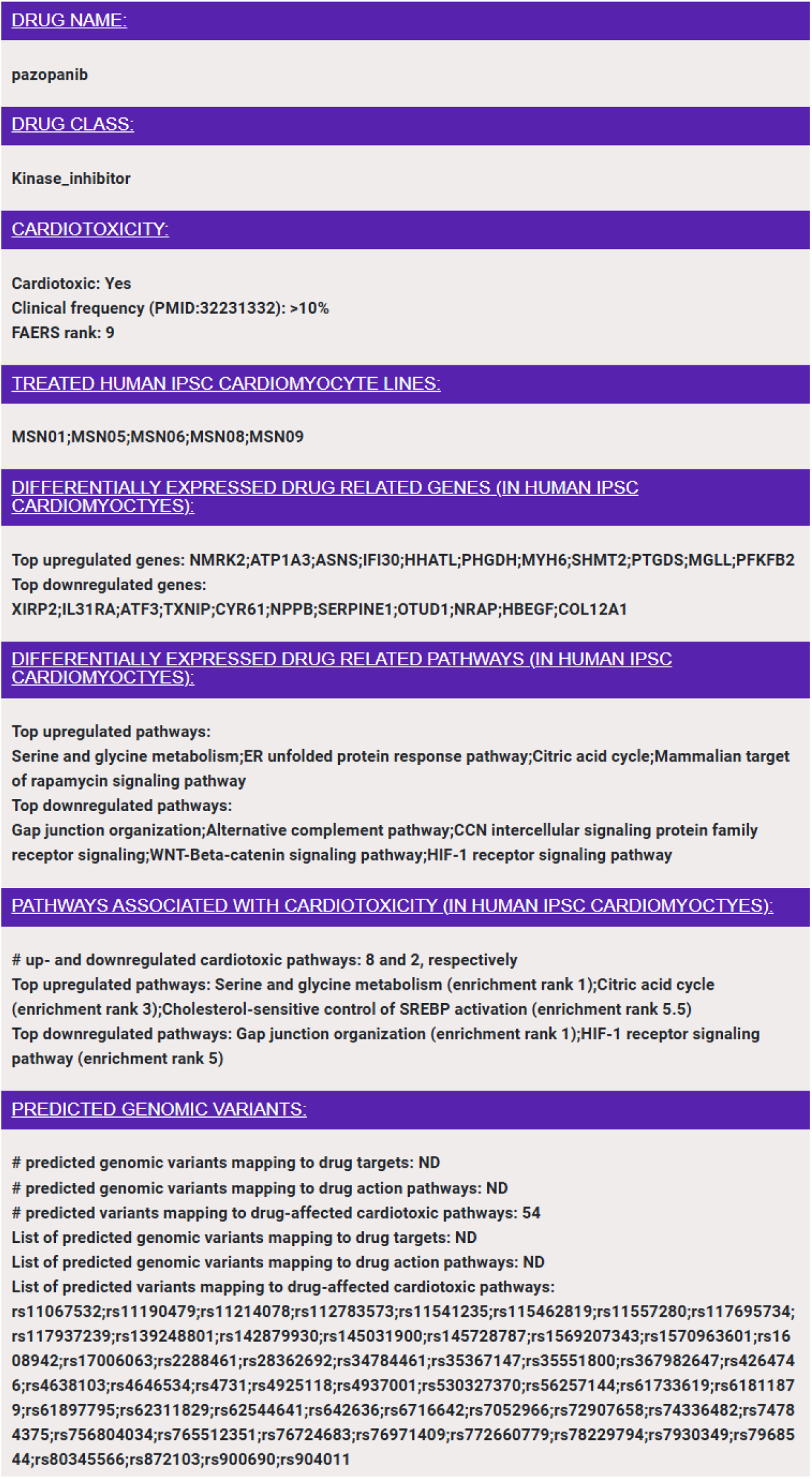
DrugTox Summary Card for pazopanib. DrugTox summary cards provide a concise summary of the drug class, its cardiotoxic potential from FAERS ranking and literature summary, ranked list of genes and pathways affected by the drug in the indicated human IPSC cardiomyocyte lines and potential genomic variants.

### Drug-induced pathways associated with cardiotoxicity

Pathways associated with cardiotoxicity for multiple drugs in multiple iPSC-derived cardiomyocyte cell lines can be readily obtained by clicking on the “Pathways” button in the “Summary of Signatures for Cardiotoxicity” window on the home page, as described above. This page gives a list of the top 25 ranked pathways associated with cardiotoxic drugs and the associated direction of change (i.e., up- or downregulation) (Fig. 3A), as well as a listing of all of genes that belong to the affected pathway. More detailed information including the identity of the drugs that regulate a pathway of interest in which iPSC-derived cardiomyocyte cell lines (Fig. 3B) can be obtained by clicking on “Drug-induced pathways associated with Cardiotoxicity” in the section entitled “Bulk Transcriptomic Datasets - Metadata”.

**Fig. 3.**
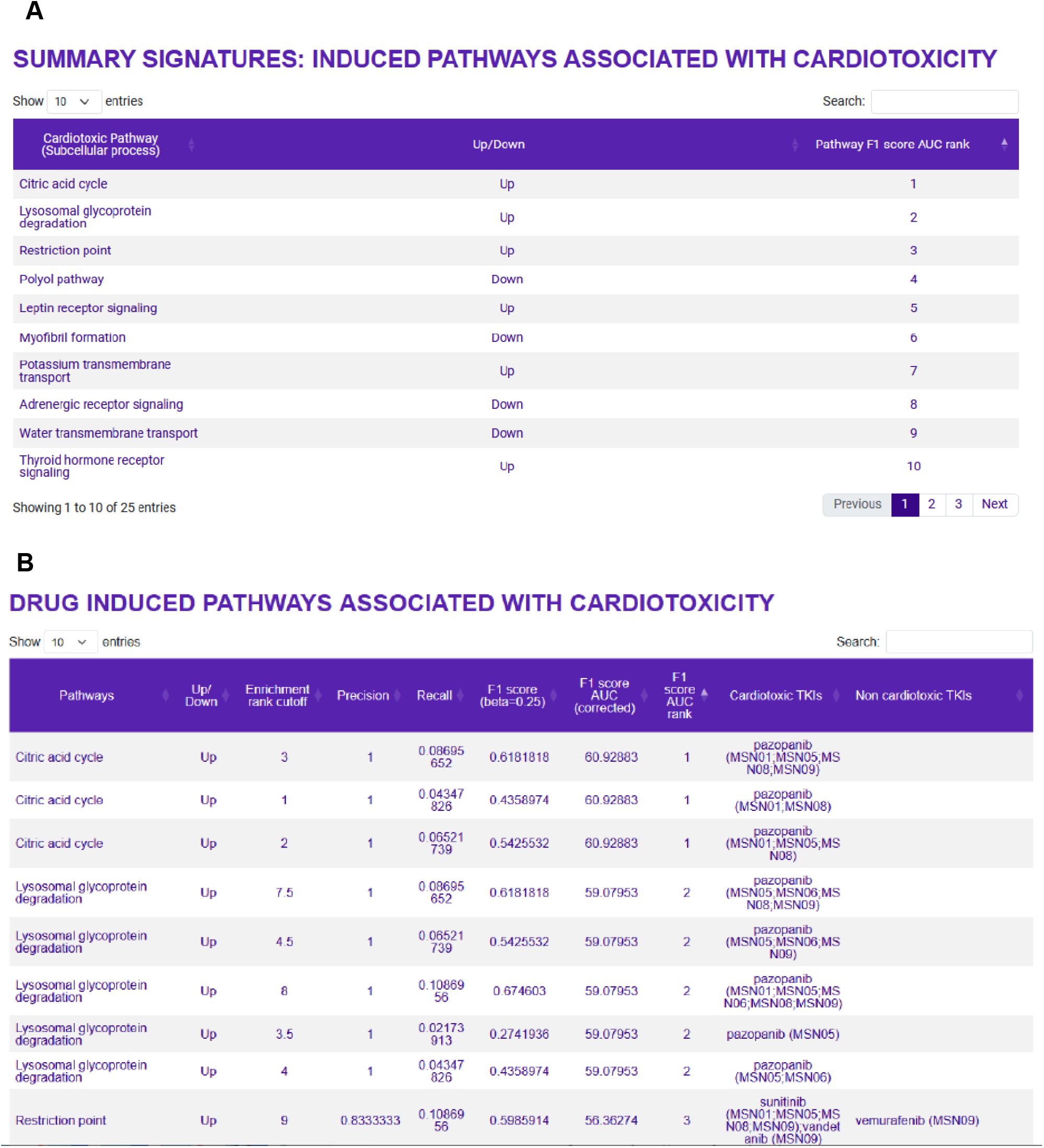
Pathways predicted to be associated with TKI-induced cardiotoxicity. **(A)** The top 25 up- and downregulated pathways that were predicted to be associated with TKI-induced cardiotoxicity can be queried on the “Drug-induced pathways associated with cardiotoxicity” page. **(B)** The second spreadsheet shows the pathway genes induced by cardiotoxic TKIs in the six human iPSC-derived cardiomyocyte cell lines used in this study.

### Evaluation of clinical drug cardiotoxicity

The “Drugs” page accessible from the “Downloadable Datasets” window provides summary information for all drugs in this database, including drug classes and their cardiotoxic potential. ROR for each drug’s cardiotoxic effect and their 95% confidence intervals were calculated using data of the FAERS, focusing on the ontology term cardiac disorder AE (OAE 0000084) as defined by the Ontology of Adverse Events ^14^. Drugs are ranked by ROR. Additionally, we extracted risk frequencies from a detailed review ^7^ summarizing the clinical literature. Risk evaluation by both methods are in good agreement. Furthermore, this page provides information on the date of approval and human drug target proteins curated from Drug Bank.^19^

### Detailed datasets

Besides providing details about the predicted pathways associated with cardiotoxicity the section entitled “Bulk Transcriptomic Datasets - Metadata” in the window “Downloadable Datasets” offers all datasets that we used to extract our summary information:

1. Drug-induced DEGs in each cell line (Download ONLY)
2. Drug-induced DEGs across all cell lines (Download ONLY)
3. Drug-induced pathways in each cell line (calculated from 1)
4. Drug-induced pathways across all cell lines (calculated from 2)
5. Induced pathways associated with cardiotoxicity
6. Predicted genomic variants influencing drug action
7. Predicted genomic variants influencing cardiotoxic drug-induced pathways

In addition, this section gives links to metadata information:

1. Cell Line Metadata
2. Experimental Metadata

In the supplementary section of our recent research publications we give detailed descriptions of the methodologies for the generation and calculation of all provided datasets.^11,23^

### Genomic Variants

The section “Bulk Transcriptomic Datasets - Metadata” provides access to datasets listing potential genomic variants that could be associated with increased (or decreased) risk for cardiotoxic drug effects. We focus on two sets of variants that could interfere with a drug’s effect on the heart: variants that map to pathways predicted to be associated with TKI cardiotoxicity or to genes coding for proteins involved in a drug’s PD or PK. For both sets we ensured that the population-wide frequency of the characterized variant alleles (≤10%) matches frequencies for cancer drug cardiotoxicity from clinical data ^7^ and that the variants are relevant for heart tissue by mapping to exon-coding regions or being part of cis-e or s-QTLs from GTEx data.^18^ It is possible that other criteria may need to be considered in future models to predict cardiotoxicity with greater accuracy. Variants mapping to the genes of pathways whose up- or downregulation is associated with cardiotoxicity are linked to a drug of interest, if that drug regulates the pathway in the same direction. We hypothesized that potential cardiotoxic variants mapping to PK/PD mechanisms should induce a deviating transcriptomic response in one of the treated cell lines. We therefore searched cell lines with transcriptomic outlier responses to a drug of interest for variant alleles that meet our population-wide and relevance criteria, documented higher counts in the outlier than in all other five cell lines and mapped to genes involved in PK/PD mechanisms of the drug of interest. The page “Potential genomic variants influencing drug PK or PD” provides the results of our predictions (Fig. 4). It shows the variant ID, the gene and genomic region the variant maps to, the relation of the mapped gene to the PK/PD protein, the PK/PD protein, the drug, its cardiotoxicity status and drug class. As indicated by the relationship category, variants can either map directly to drug target proteins, transporters or metabolizing enzymes or to transcription factors and kinases potentially regulating abundances and activity levels of those proteins.

**Fig. 4.**
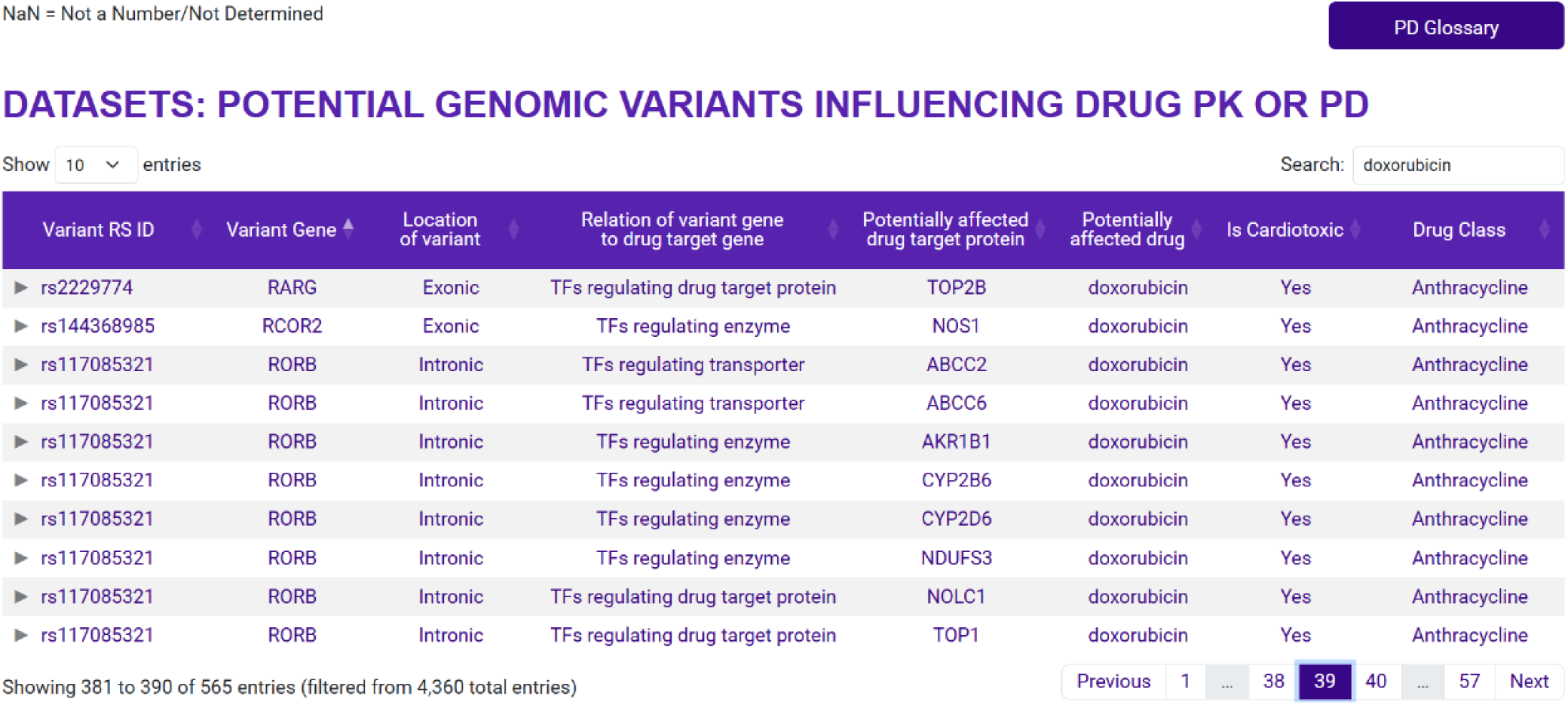
Genomic variants that are potentially associated with a drug’s cardiotoxicity by interfering with the drug’s PD or PK. Note that some genomic variants can map to multiple genes.

### Multicompartment ODE models for numerical simulations

Since the physiological effects of drugs are time dependent, multicompartment ODE models are widely used in PK/PD studies. In connecting the transcriptomic data to numerical simulations, frequently, we have to make assumptions regarding proportionality between changes in the levels of transcripts to changes in protein and activity levels. Making such assumptions we provide two models, one for arrhythmia and another for cardiomyocyte hypertrophy. These models can serve as a starting point for detailed computational modeling of cardiotoxic effects of drugs. Downloaded MATLAB code can be run without any further interventions to simulate the drug effects. The associated ReadMe files and the published references give useful advice for running the numerical simulations.

### Use Case

predicTox-KE presents multiple genomic variant candidates that could be associated with an increased (or decreased) risk for development of cardiac drug side effects. As described above, to influence a drug’s cardiotoxic potential a variant could map to a gene that participates in a pathway whose altered expression is associated with cardiotoxicity or map to a gene whose gene product participates in a drug’s PK or PD. predicTox-KE provides variant candidates for both groups. Here we focus on a variant associated with PD response. Using our algorithm and without using the genome-wide association study (GWAS) data, we found the variant rs2229774 in the exon coding region of RARG in the one cardiomyocyte cell line that shows an outlier transcriptomic response to anthracycline treatment ^11^. This variant is one of three variants identified in GWAS with the strongest evidence for anthracycline-induced cardiotoxicity (AIC) ^24^. It is casually linked to AIC, since it lies within the coding region of the transcription factor gene RARG that regulates the expression of the anthracycline target protein DNA topoisomerase TOP2B.^25^ The mechanism of action for the variant identified by our computational analysis indicates this connection as well (Fig. 4).

As our initial results show, the strategy we use can identify additional genomic variants associated with cardiac safety. This example demonstrates how predicTox can be used to identify potential genomic variants for target GWAS with lower sample size requirements or experimental testing for drug safety in humans. As has been done for anthracycline toxicity, ^25^ gene editing studies in iPSC-derived cardiomyocytes could constitute a potential experimental test for a predicted variant. To obtain candidate lists of genomic variants and associated genes select the “Bulk Transcriptomic Datasets-Metadata” button in the predicTox homepage, as described above. Then select “Predicted genomic variants influencing drug action” in the white “Datasets” window. Rows can be sorted by “Potentially affected drug” or any other entries. To focus on variants mapping to coding regions, which should be the most likely candidates for monogenic traits ^26,27^ enter “exonic” into the “Search” field. To investigate which drugs, genes and PD/PK proteins might be linked to that variant, copy-paste its reference single nucleotide polymorphism ID (rsID) into the search field.

## Conclusion

The expansion of data in different domains including genomics, cell biology of drug action and transcriptomic profiles offer new data integration opportunities that produce synergistic new knowledge. The new knowledge can be accessed by from knowledge environments such as predicTox. With the complementary development of human iPSC-based cell systems and organoids for cardiac tissues,^28^ new cell-based methods that can reduce animal testing ^29^ for discovery of cardiac drugs and for cardiac safety testing are becoming realistic options. Beyond cardiotoxicity the data integration approaches we use here can be readily used to understand and predict genomic variants for hepatotoxicity, nephrotoxicity and pulmonary toxicity, both common and rare. Databases such as predicTox KE that are centered around integration of data from different domains can play a useful role in using such data integration and modeling to drive the design of human cell-based experiments for drug safety.

## Funding

predicTox was developed with support from the United States FDA (FDA BAA Contract No. 75F40119C10021). All transcriptomics data were obtained and analyzed by the LINCS DToxS Center funded by the NIH Common Fund LINCS Program (U54HG 008098).

## Conflicts of Interest

None.

## Disclaimer

The article reflects the views of the authors and should not be construed to represent the FDA’s views or policies.

## Notes

https://predictox.org/

### Competing Interest Statement

The authors have declared no competing interest.

